# LIMPACAT : Multi-Omics Attention Transformer for Immune Prediction in Liver Cancer Using Whole-Slide Imaging

**DOI:** 10.1101/2024.10.28.620410

**Authors:** Yen-Jung Chiu

## Abstract

This study introduces LIMPACAT (Liver Immune Microenvironment Prediction and Classification Attention Transformer), a framework that leverages whole-slide images (WSIs) to predict immune cell levels associated with prognosis in liver cancer. Since direct immune cell composition data is unavailable in the TCGA-LIHC dataset, we inferred immune cell levels using liver-specific single-cell RNA sequencing (scRNA-seq) data to simulate bulk RNA-seq profiles, allowing estimation of immune compositions within the tumor microenvironment. To ensure consistency, we tested three normalization methods—log normalization, canonical correlation analysis (CCA), and SCTransform—for scRNA-seq data integration and preprocessing, which facilitated reliable predictions of immune cell distributions in bulk RNA-seq.

By applying this scRNA-seq-informed cell composition deconvolution model to real bulk RNA-seq data from liver cancer samples, we confirmed alignment in immune cell composition estimates between bulk RNA-seq and scRNA-seq. Using a multiple instance learning (MIL) framework with an attention transformer, LIMPACAT achieved approximately 80% accuracy in classifying immune cell levels relevant to patient prognosis. These findings highlight the feasibility of integrating WSIs and multi-omics data to enhance immune profiling and prognostic predictions in liver cancer research.

## Background

The tumor microenvironment (TME) is a complex biological system composed of not only cancer cells but also a variety of immune cells. Interactions between immune cells and cancer cells within the TME have been shown to influence cancer growth and metastasis^1^. For instance, high levels of memory CD8+ T cells are associated with better patient outcomes^2^. However, upon interaction with cancer cells, CD8+ T cells may transform into an “exhausted” state, which is linked to poorer prognosis in patients exhibiting high exhausted CD8+ T cell expression^3^. Additionally, differences in immune cell subtypes, characterized by distinct surface proteins, may lead to opposite effects on cancer cells. For example, chimeric antigen receptor (CAR) T cell therapy isolates T cells from the body, activates them to attack cancer cells, but these T cells can still be restricted by the immunosuppressive TME. Regulatory T cells (Tregs), for instance, suppress immune responses, reducing the efficacy of CAR T cell therapy^4^. Similarly, immune checkpoint inhibitors (ICIs) are a type of cancer therapy that blocks proteins such as PD-1, PD-L1, and CTLA-4, which otherwise inhibit immune cell function and contribute to T cell exhaustion, thereby increasing the ability of immune cells to attack cancer cells^5^. This evidence highlights the importance of analyzing the immune TME to improve cancer treatment and prognosis. Although flow cytometry can also determine immune cell types, it requires a large number of live cells and is limited in its ability to analyze cancer tissues^6^. The CCD algorithm leverages computational biology, facilitating data reuse and large-scale data analysis to assess TME influence on cancer^7–10^. Rather than replacing biological tests, CCD algorithms identify key biological features from existing data, aiding researchers in uncovering potential insights for cancer exploration.

In recent years, the availability of single-cell RNA sequencing (scRNA-seq) data has increased significantly through public databases such as NCBI GEO and 10x Genomics. Compared to bulk RNA-seq or microarray data, scRNA-seq provides higher precision by analyzing individual cells, enabling researchers to identify immune cell subtypes and their interactions within sequencing samples, such as exhausted CD8+ T cells and FCGR3A+ macrophages in liver cancer^11^. However, scRNA-seq data presents challenges with batch effects, where data integration from diverse sources is difficult without normalization. To address this issue, various normalization techniques, such as log normalization, canonical correlation analysis (CCA), and scttransform, have been developed to mitigate batch effects, allowing more accurate and consistent cross-sample comparisons. Effective normalization is essential for ensuring the reliability of downstream analyses, particularly in studies investigating complex interactions within the tumor microenvironment.

Recent advancements in attention-based deep learning models have enabled significant progress in computational pathology, especially for whole-slide image (WSI) analysis. The Clustering-Constrained Attention Multiple Instance Learning (CLAM) model utilizes attention mechanisms to identify diagnostically relevant regions within WSIs under weak supervision. By assigning varying attention weights, CLAM offers enhanced interpretability and data efficiency, which is particularly beneficial in scenarios with limited annotations. However, CLAM may be constrained in capturing finer inter-regional relationships, which limits its applicability beyond classification tasks^12^. Similarly, the use of attention-based convolutional neural networks for meningioma classification has shown promise by focusing on tumor regions deemed critical by pathologists, leading to accurate molecular classification. Although beneficial for accelerating precision medicine workflows, this approach requires extensive management of multi-layered attention and may be challenging to generalize to other tumor types^13^. Meanwhile, the MHAttnSurv model for survival prediction employs multi-head attention to explore diverse morphological patterns across tumor slides, enhancing the robustness of survival prediction. While effective, MHAttnSurv demands high computational resources, posing challenges for large WSI applications^14^.

Histopathological image digital pathology is a critical technique in cancer diagnosis, offering a simpler and more cost-effective alternative to molecular experiments^15^. Hematoxylin and eosin (H&E) staining is widely used in digital pathology, providing contrast among various tissues by staining cytoplasm^16^, cell nuclei, extracellular matrix, and other cellular structures^15,17^. These histopathological findings can correlate with prognosis; however, manual interpretation of large-scale pathological images is impractical. Known variations in immune cell abundance also impact patient outcomes, emphasizing the need to accurately classify immune cell levels through digital pathology. To address this, we developed LIMPACAT (Liver Immune Microenvironment Prediction and Classification Attention Transformer), a model designed to predict immune cell abundance associated with liver cancer prognosis directly from pathological images. By facilitating a deeper understanding of immune responses in the tumor microenvironment, LIMPACAT supports prognostic assessments and advances the capabilities of digital pathology.

## Methods

### Data Download and Pre-processing

The scRNA-seq data in this study is derived from the GSE189903 dataset, initially containing 34 primary liver cancer samples, including hepatocellular carcinoma (HCC) and intrahepatic cholangiocarcinoma samples from various tumor regions^18^. For the analysis, 20 HCC samples were selected, each divided into distinct expression regions. The data processing pipeline, implemented with the Seurat R package^19^, included quality control (QC) steps to filter out low-quality cells based on gene count and mitochondrial content. After QC, three normalization methods— lognormalization^20^, canonical correlation analysis (CCA)^21^, and scttransform^20^—were applied to correct for batch effects, ensuring consistency across samples. This preprocessed scRNA-seq data served as a foundation for accurately predicting immune cell abundance associated with liver cancer prognosis.

Liver cancer data was downloaded from the TCGA-LIHC dataset available on the Genomic Data Commons Data Portal. This dataset includes gene expression and diagnostic imaging data along with clinical information. Gene expression data was preprocessed using the UQ normalization method.

### Evaluation of scRNA-seq Normalization

To evaluate the clustering consistency and the effectiveness of batch effect correction, we performed adjusted rand index (ARI) comparisons across different conditions^22–25^. The ARI, calculated as

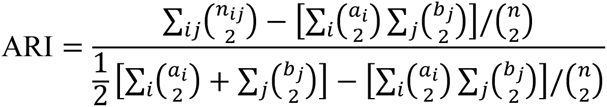

measures the alignment between clustering results and reference groupings, adjusting for chance alignment. Here, *n* represents the total number of data points (e.g., cells), *n_ij_* is the number of data points shared between cluster *i* in one clustering and cluster j in the reference clustering. The sums *a_i_* and *b_j_* represent the total number of points in cluster *i* and cluster *j*, respectively, for each clustering method. We assessed the alignment between cell type annotations, clustering results, and liver region-specific batch effects, which were derived from multiple patients and liver regions within the dataset.

1. A-C (Annotation vs. Clustering): Cell annotation was performed using SingleR, which classified cells into biologically meaningful types based on reference datasets. Clustering was done using Seurat’s unsupervised method. ARI was calculated to assess the alignment between SingleR’s annotations and Seurat’s clustering. A high ARI score indicates that Seurat’s clustering successfully identifies biologically distinct cell subpopulations.
2. B-C (Batch vs. Clustering): To evaluate the effect of liver region-specific batch effects on clustering, ARI was computed between liver region batches and Seurat’s clustering results. A low ARI score is desirable here, suggesting that clustering is driven by biological signals rather than technical batch effects, indicating successful batch effect correction.
3. A-B (Annotation vs. Batch): ARI was also calculated between cell annotations from SingleR and liver region batches to examine the distribution of annotated cell types across different regions. A low ARI score in this comparison indicates that annotated cell types are consistently distributed across regions without being influenced by batch effects, reinforcing the stability of the annotations across spatially distinct liver regions.

### Immune Cell Prediction Model Development Using scRNA-seq

Cell types were annotated using the SingleR package^26^, which matched scRNA-seq data with reference profiles to identify specific immune cell types. Based on these annotations, we developed a deep learning model, ensemble deep neural network (ensemble-DNN), to predict immune cell composition from gene expression data, focusing on accurately estimating proportions of immune subtypes relevant to the tumor microenvironment.

To generate the training data for the deep learning model, we assumed that the tissue comprises a mixture of individual cells to generate the training data for the deep learning model. Based on this assumption, we constructed the RNA-seq gene expression profile of liver cancer tissue using scRNA-seq data. Specifically, we randomly sampled cells from different cell types within the liver cancer scRNA-seq dataset and aggregated their gene expression profiles. The aggregation was performed assuming that the bulk RNA-seq expression is a sum of individual cells from different types.

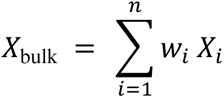

Where:

- *X*_bulk_ represents the gene expression profile for the simulated bulk RNA-seq sample.
- *X*_*i*_ is the gene expression profile of the *i-th* cell type.
- *w*_*i*_ is the weight or proportion of the *i-th* cell type in the simulated tissue, where 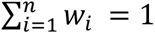.
- *n* is the total number of cell types.

### Digital Pathology Image Analysis

In this study, digital pathology analysis was conducted using WSIs from H&E. These WSIs are rich in spatial information and provide a comprehensive view of tissue morphology. The objective was to leverage these images for weakly supervised learning in the context of immune cell type prediction and abundance estimation within the TME.

We employed two parallel approaches for model development: (1) using the CLAM toolkit, a weakly supervised learning framework optimized for feature extraction from high-resolution images; and (2) building a custom deep learning workflow using python package MONAI, a medical imaging toolkit that allows greater flexibility in designing general CNN-based, attention-based (ATT), and transformer attention-based (ATT_TRANS) architectures in multiple instance learning.

## Results

### Study Design

This workflow is designed to investigate the potential immune cell compositions within bulk RNA-seq that may be associated with patient prognosis through the analysis of scRNA-seq data. Additionally, by employing WSI analysis, the model aims to predict immune cell levels and ultimately forecast patient prognosis based on these WSIs (Figure 1). Initially, we performed datasets download and filtering to obtain scRNA-seq datasets, including both malignant and non-malignant samples, providing a comprehensive view of the liver tumor microenvironment. Rigorous quality control procedures were applied at this stage, eliminating low-quality cells and retaining only those with acceptable feature counts and mitochondrial gene proportions.

**Figure 1.**
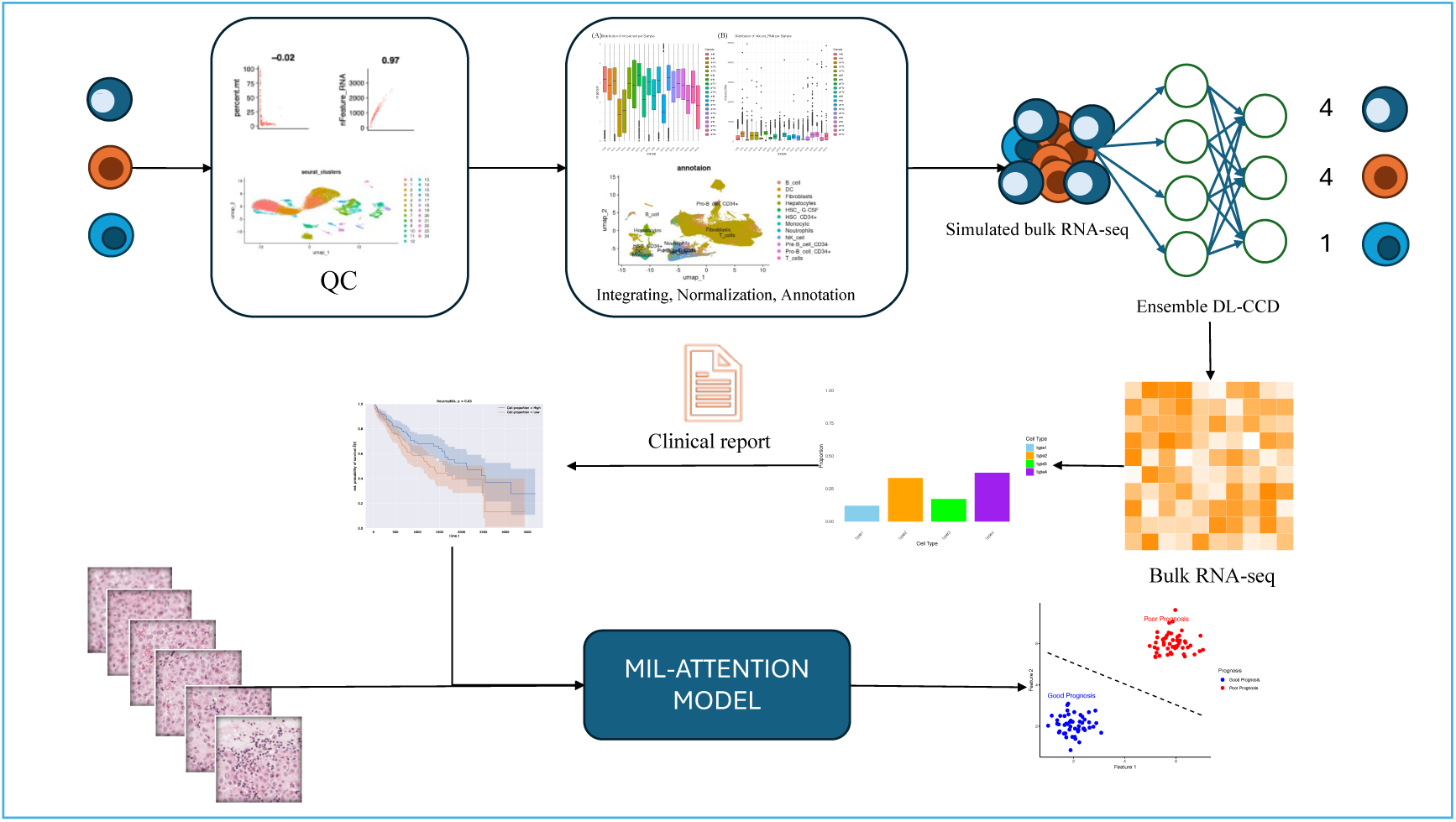
Workflow of LIMPACAT

Following QC, samples integration and normalization were conducted to combine data from multiple sources and reduce batch effects across samples. Seurat’s normalization methods were applied, and various approaches were tested, including lognormalization, CCA, and scttransform. These methods were evaluated for their effectiveness in mitigating batch variability, enabling consistent cross-sample comparisons. After normalization, clustering and cell type annotation was performed using dimensionality reduction techniques, specifically Uniform Manifold Approximation and Projection (UMAP), to group cells based on gene expression profiles. This clustering step facilitated the identification of cell populations with similar transcriptional signatures, delineating distinct clusters corresponding to specific cell types or functional states within the tumor microenvironment. Subsequently, cell type annotations were assigned by examining characteristic gene expression patterns within each cluster.

Next, we assessed the effectiveness of each normalization method using the ARI, a metric for clustering consistency. This evaluation was crucial to confirm that the selected normalization approach optimized clustering accuracy, supporting reliable downstream scRNA-seq analyses in the liver cancer research framework. And, we created simulated bulk RNA-seq datasets based on scRNA-seq data using a random sampling method and developed a cell composition deconvolution model using an ensemble deep learning framework. This model was used to predict immune cell content in real bulk RNA-seq data, and these predicted values were analyzed in survival studies to identify immune cells associated with patient prognosis. Last step, survival analysis groups were used to train a multiple instance learning (MIL) attention model, enabling WSI image analysis to predict immune cell abundance. This immune cell abundance, directly related to survival analysis, allows for predictions of patient outcomes based on immune cell levels.

### Gene Count and Quality Control Metrics of the GSE189903 Dataset

The GSE189903 dataset provides a scRNA-seq dataset for 20 liver cancer samples, encompassing approximately 737,280 cells in total. The gene count within each dataset ranges from 17,000 to 20,000, with an overlapping set of 11,632 genes shared across samples. This extensive coverage allows for a broad view of gene expression, offering insights into the diversity of cell populations within the liver tumor microenvironment.

The data filtering process for the GSE189903 dataset highlights the variation in gene counts per cell before filtering, ranging from approximately 12,000 to over 20,000 genes. This wide range reflects the inherent diversity in gene expression among cells, influenced by both biological variability and potential technical noise (Figure 2 A). After quality control, the number of retained cells varies across samples, with some retaining close to 15,000 cells, while others have fewer than 5,000, suggesting differences in cell quality across samples (Figure 2 B). Most samples maintain between 15,000 and 20,000 genes post-filtering, which preserves a substantial level of gene expression diversity (Figure 2 C). Additionally, the proportion of retained cells per sample generally ranges from 5% to 25% of the original count, indicating the effectiveness of stringent quality control in removing low-quality cells while retaining adequate data diversity for analysis (Figure 2 D). The correlation between various filtering metrics, including Raw Features, Filtered Cells, Filtered Features, and the percentage of retained cells, is illustrated in the correlation matrix (Supplementary Figure 1). This matrix shows positive correlations between these metrics, reinforcing the consistency and impact of the filtering criteria in selecting high-quality cells and features for downstream analyses.

**Figure 2.**
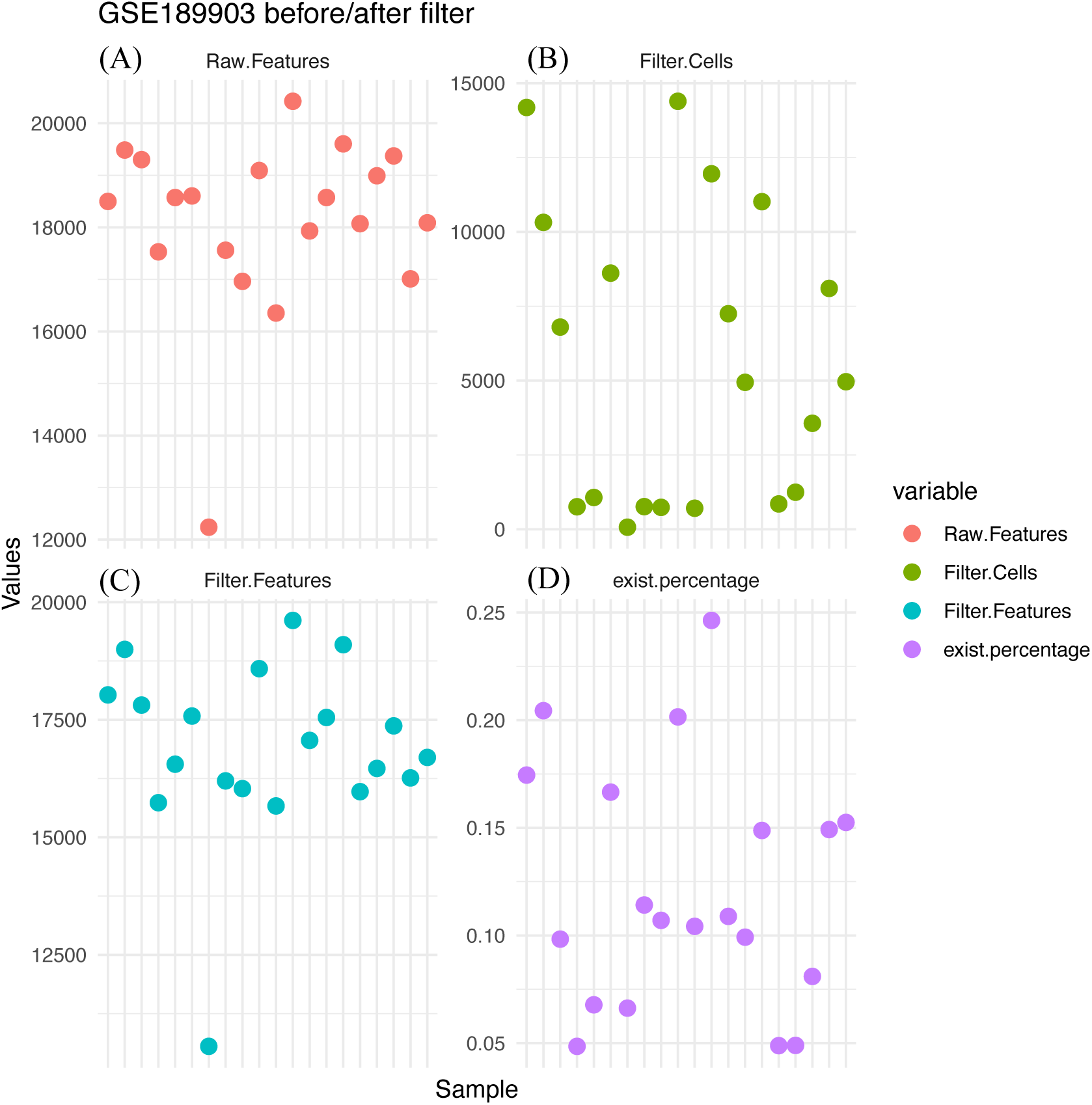
Distribution of Features and Cells Before and After Filtering in GSE189903 Dataset

### Correlation Analysis and Data Quality Evaluation

The distribution of nFeature across the dataset further underscores the range of detected genes per cell, with most cells containing fewer than 100 genes, though a significant subset holds between 250 and 2,500 genes (Supplementary Figure 2). After data filtering, each sample’s mitochondrial gene content (mt percent) and nCount distributions were assessed to evaluate cell quality and sequencing depth. The mt percentage statistics across samples show variability in mitochondrial gene content, with mean values ranging from approximately 1.42 to 3.38. Median mt percentages are generally close to the mean values, suggesting a relatively symmetric distribution within samples. Standard deviations range from 0.85 to 1.38, indicating moderate dispersion around the mean. Minimum mt percentages are consistently 0 across samples, while maximum values are near 5, reflecting the upper limit set by the filtering criteria. Cell counts per sample vary from a low of 69 cells in 2HN to a high of 14,183 cells in 1HB, illustrating differences in data retention across samples following filtering (Supplementary Figure 3A and Supplementary Table 1). The nCount distribution across samples reveals significant variability in sequencing depth, with mean values ranging from approximately 879 to 4204. Median nCounts vary similarly, indicating differences in sequencing coverage across cells in each sample. The standard deviation values reflect considerable dispersion within samples, with certain samples displaying a wider range of nCounts, from minimums around 280 to maximums exceeding 40,000. Cell counts for each sample also vary, from a low of 69 cells in sample 2HN to a high of 14,183 in sample 1HB, underscoring the diversity and heterogeneity in data quality and cell characteristics across the dataset (Supplementary Figure 3B and Supplementary Table 2). Cells were filtered based on two primary criteria to ensure high-quality data. First, cells with nFeature counts between 250 and 2,500 were retained to exclude potential sequencing artifacts or low-quality cells. Second, cells with mitochondrial gene percentages exceeding 5% were removed, as elevated mitochondrial content generally correlates with dying or dead cells. This filtering process retained approximately 10-25% of the original cells, with counts ranging from several dozen to tens of thousands across samples (Supplementary Table 3). Moreover, correlation analysis was performed to assess further data quality and the potential impact of technical variation. Analysis of nCount against mitochondrial percentage yielded a weak correlation, suggesting that mitochondrial gene content was independent of sequencing depth (Supplementary Figure 4). Conversely, nCount and nFeature exhibited a strong positive correlation, reinforcing the reliability of the retained cells after filtering (Supplementary Figure 5).

### Comparison of scRNA-seq Normalization Methods

To integrate data from multiple sample sources and eliminate batch effects, we compared three normalization methods: log normalization, CCA, and scttransform. Each method was evaluated based on clustering consistency and sample distribution using UMAP and ARI.

For log normalization, UMAP clustering resulted in the definition of 25 clusters (Figure 3 A). When comparing the consistency of clustering between 20 sample types and the 25 clusters generated by log normalization, an ARI score of 0.168 was achieved (Figure 3 B). In contrast, CCA normalization produced 28 clusters (Supplementary Figure 6 A) with an ARI score of 0.04 (Supplementary Figure 6 B), which is considerably lower than that achieved with log normalization. Scttransform normalization generated 23 clusters (Supplementary Figure 7 A), with an ARI score of 0.043 (Supplementary Figure 7 B). This method showed an intermediate performance between log normalization and CCA. Log normalization achieved the highest ARI score in most comparisons, marked in yellow, indicating its superior performance in clustering consistency (Supplementary Table 4).

**Figure 3.**
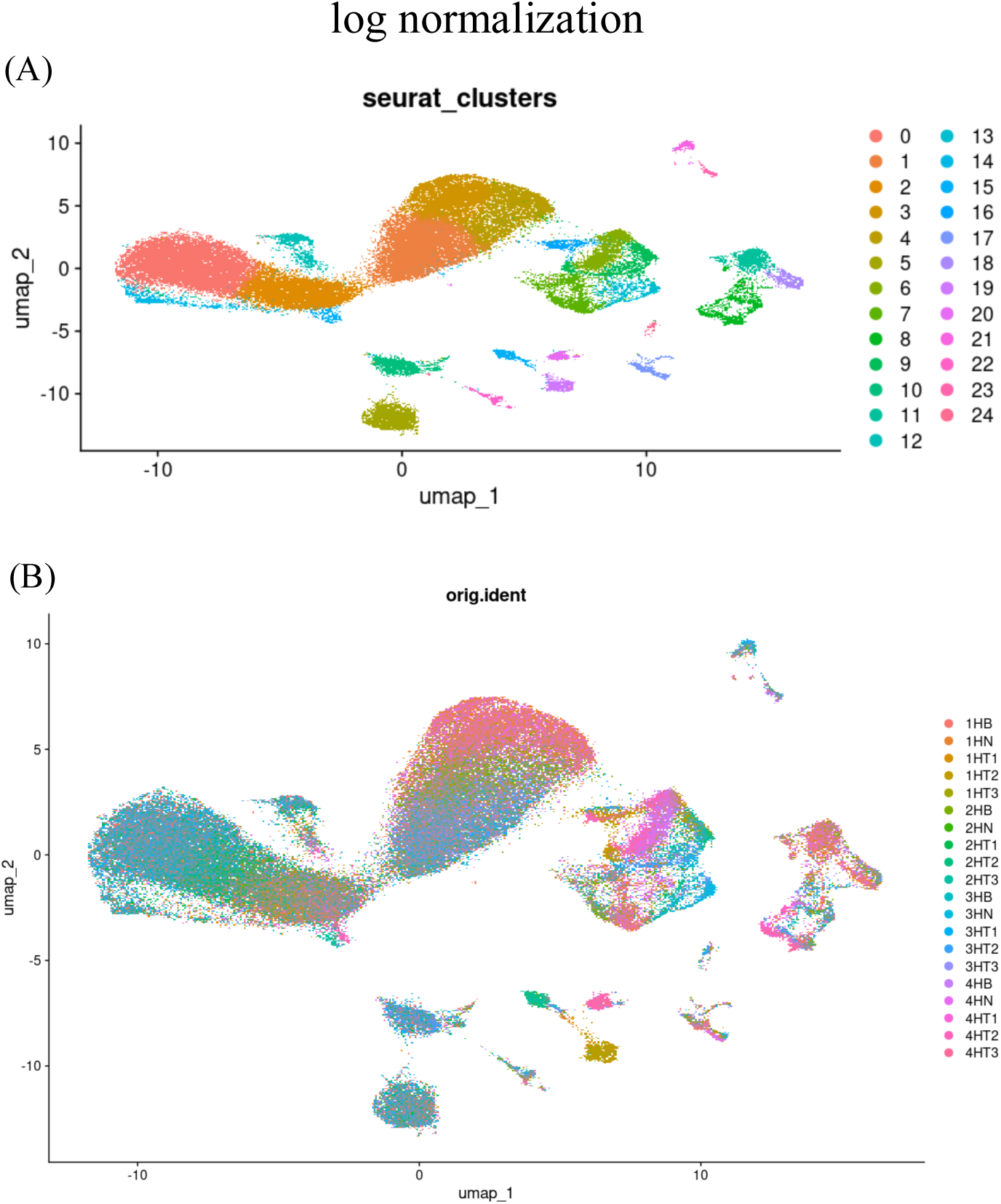
UMAP clustering of scRNA-seq data post-log normalization. (A) UMAP plot showing 25 Seurat-defined clusters, capturing cellular diversity within the dataset. (B) UMAP plot with cells colored by sample identity, illustrating the distribution of cells across clusters from different samples.

### Comparison of Clusters Correlation

To assess the impact of log normalization, CCA, and SCTransform on data structure, we examined the nFeature and nCount distributions across samples, as well as sample-to-sample correlation patterns.

For log normalization, the nFeature and nCount distributions showed relatively wide ranges across samples, reflecting a preserved heterogeneity in gene expression levels (Figure 4 A). This variation suggests that log normalization maintains biological diversity across cell populations, although some technical noise may remain, as evidenced by the dispersed distributions in certain samples. Additionally, the sample-to-sample correlation heatmap revealed moderate correlation consistency across samples, with notable variability (Figure 4 B), indicating partial alignment of gene expression profiles but potential residual batch effects.

**Figure 4.**
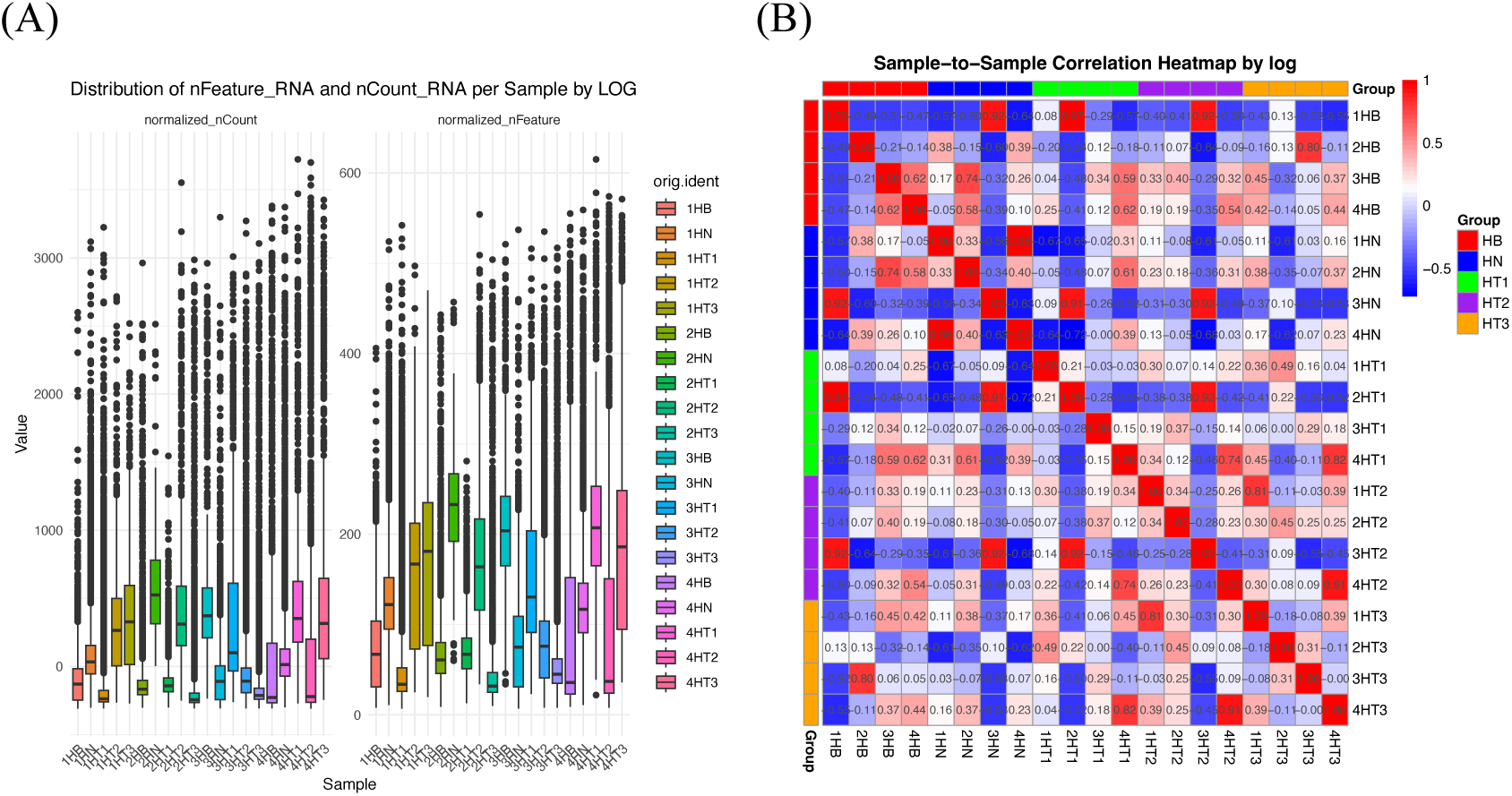
nFeature and nCount distributions (A) and sample-to-sample correlation heatmap (B) for log normalization. The boxplots show a balanced distribution across samples, while the heatmap indicates high consistency.

In contrast, CCA normalization produced more concentrated nFeature and nCount distributions, showing a narrower range of values across samples (Supplementary Figure 8 A). This consistency suggests that CCA effectively reduces batch effects, providing a high level of standardization and data alignment. The sample-to-sample correlation heatmap for CCA normalization demonstrated strong correlations among samples, indicating robust data integration and minimal batch variability (Supplementary Figure 8 C). The concentrated distributions and high correlation values suggest that CCA achieves a significant level of data harmonization across samples.

With sct normalization, the nFeature and nCount distributions were more uniform across samples than with log normalization, yet displayed a slightly broader range compared to CCA (Supplementary Figure 8 B). This intermediate spread indicates that sct normalization balances standardization with the retention of biological variability, making it suitable for analyses requiring both consistency and biological relevance. The sample-to-sample correlation heatmap for sct normalization displayed high correlations across samples, similar to CCA, indicating effective data integration with minimal technical noise (Supplementary Figure 8 D).

### Single-Cell Type Annotation

To classify cell types within liver cancer samples, we applied the SingleR package across three normalization methods (log normalization, CCA, and scttransform) to perform single-cell annotation. Each method consistently indicated a high prevalence of T cells within the dataset, followed by monocytes/NK cells and cancer cells. Conversely, neutrophils, hematopoietic stem cells (HSC), and Pre-B cells were observed in notably lower quantities across all three normalization methods (Supplementary Table 5-7). For the log normalization approach, UMAP clustering revealed the distribution of annotated cell types, highlighting the abundance of T cells (Figure 5 A). Supplementary Table 5 provides a quantitative breakdown of each cell type, where T cells accounted for the largest portion, with a total of 94,602 cells. Monocytes (3,979 cells) and NK cells (4,576 cells) were also relatively abundant, whereas neutrophils and specific HSC subtypes were present in minimal quantities. Using the CCA normalization method, UMAP visualization identified 12 distinct cell types, with T cells remaining the most prominent cell type, totaling 97,005 cells (Supplementary Figure 9 A). As summarized in Supplementary Table 6, monocytes (3,299 cells) and NK cells (2,615 cells) followed in frequency, while neutrophils and HSC subtypes again demonstrated low representation. The scttransform normalization approach yielded comparable findings, with T cells as the predominant cell type (96,693 cells) according to UMAP analysis (Supplementary Figure 9 A). Supplementary Table 7 details the cell counts, where monocytes (4,351 cells) and NK cells (2,564 cells) maintained similar distributions as seen in previous methods. Neutrophils and HSCs were again sparse, underscoring a consistent trend across normalization methods. A comparative overview of the cell type distributions across the three normalization methods is presented in Figure 5 B. These results highlight the robust nature of the SingleR package in identifying major immune cell types within liver cancer samples, as well as its consistency in detecting rare cell types, such as neutrophils and specific HSC subtypes, across various normalization techniques.

**Figure 5.**
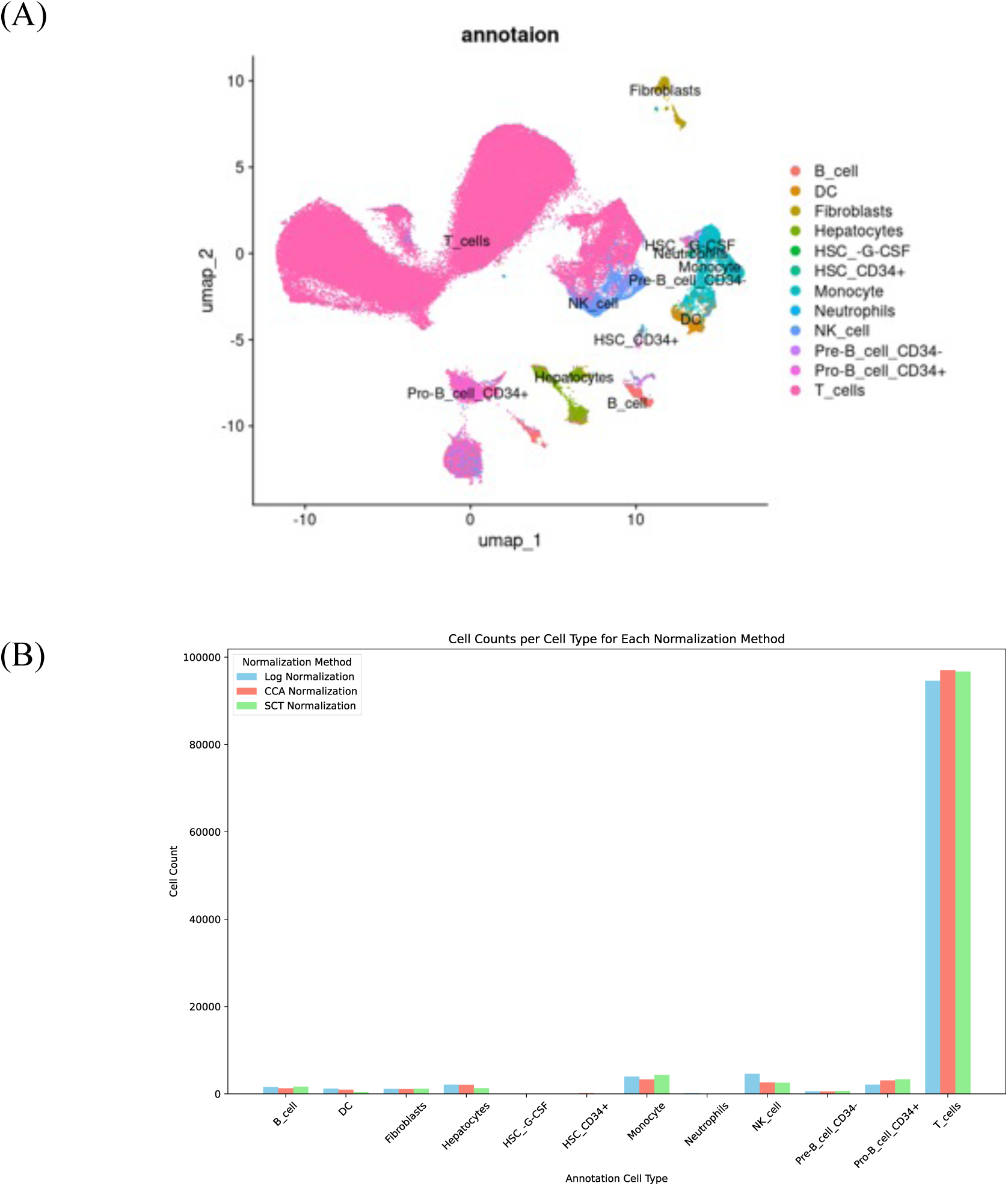
(A) shows the UMAP visualization of cell type annotations following log normalization, illustrating the distribution of major immune cell types, such as T cells and monocytes, along with rarer cell types like neutrophils and specific HSC subtypes. (B) provides a bar chart comparing cell type distributions across the three normalization methods (log, CCA, and SCT).

### Cell Composition Deconvolution Model and Immune Cell Level Estimated from WSI

To develop a model for analyzing digital pathology slides and predicting immune cell levels associated with patient prognosis, this study utilized the TCGA-LIHC dataset. As the TCGA-LIHC dataset does not provide direct information on immune cell composition, the first step was to infer immune cell composition for each liver cancer sample. Using liver-specific scRNA-seq data and applying three different normalization methods, simulated bulk RNA-seq expression profiles were generated, with immune cell counts as markers. The cell deconvolution model using ensemble-DNN, was trained with 8,000 simulated datasets and validated with 2,000 datasets, showing no signs of overfitting in accuracy and error rate (Supplementary Figure 10 A).

We then applied the developed liver scRNA-seq cell composition deconvolution model to estimate the cell composition of real bulk RNA-seq profiles from liver cancer samples. Among the three normalization methods, CCA and SCT normalized data showed similar immune cell composition distributions for LIHC samples. After consolidating the ARI and correlation results (Supplementary Figure 10 B-D and Supplementary Table 4), we selected the LOG-normalized CCD model’s predicted cell composition for survival analysis. Survival analysis indicates a positive association between higher B cells, NK cells, and monocytes and longer survival times. In contrast, a negative association is observed with elevated levels of CD34+ B cells (Supplementary Figure 10 E-H, Supplementary Table 8).

To evaluate the performance of both CLAM and our models (CNN, ATT, and ATT_TRANS) on B cell, CD34+ B cell, monocyte, and NK cell classification tasks, we stratified groups based on high and low CCD prediction scores. Statistical analysis using the log-rank test confirmed significant differences between these groups. The CLAM model achieved a baseline validation accuracy of 50% for B cell, NK cell, monocyte, and CD34+ B cell classifications. In contrast, our models demonstrated notably higher validation accuracies.

For B cell classification, CNN achieved 64%, ATT reached 72%, and ATT_TRANS obtained 73% in validation accuracy. NK cell classification validation accuracies forCNN, ATT, and ATT_TRANS were 60%, 70%, and 73%, respectively. Monocyte classification showed validation accuracies of 64% for CNN, 79% for ATT, and 76% for ATT_TRANS. Lastly, CD34+ B cell classification yielded validation accuracies of 73% for CNN, 76% for ATT, and 79% for ATT_TRANS.

These results indicate that attention-based models, particularly the transformer-enhanced ATT_TRANS, offer superior feature extraction and classification capabilities, effectively distinguishing between good and poor survival groups (Table 1).

**Table 1.**
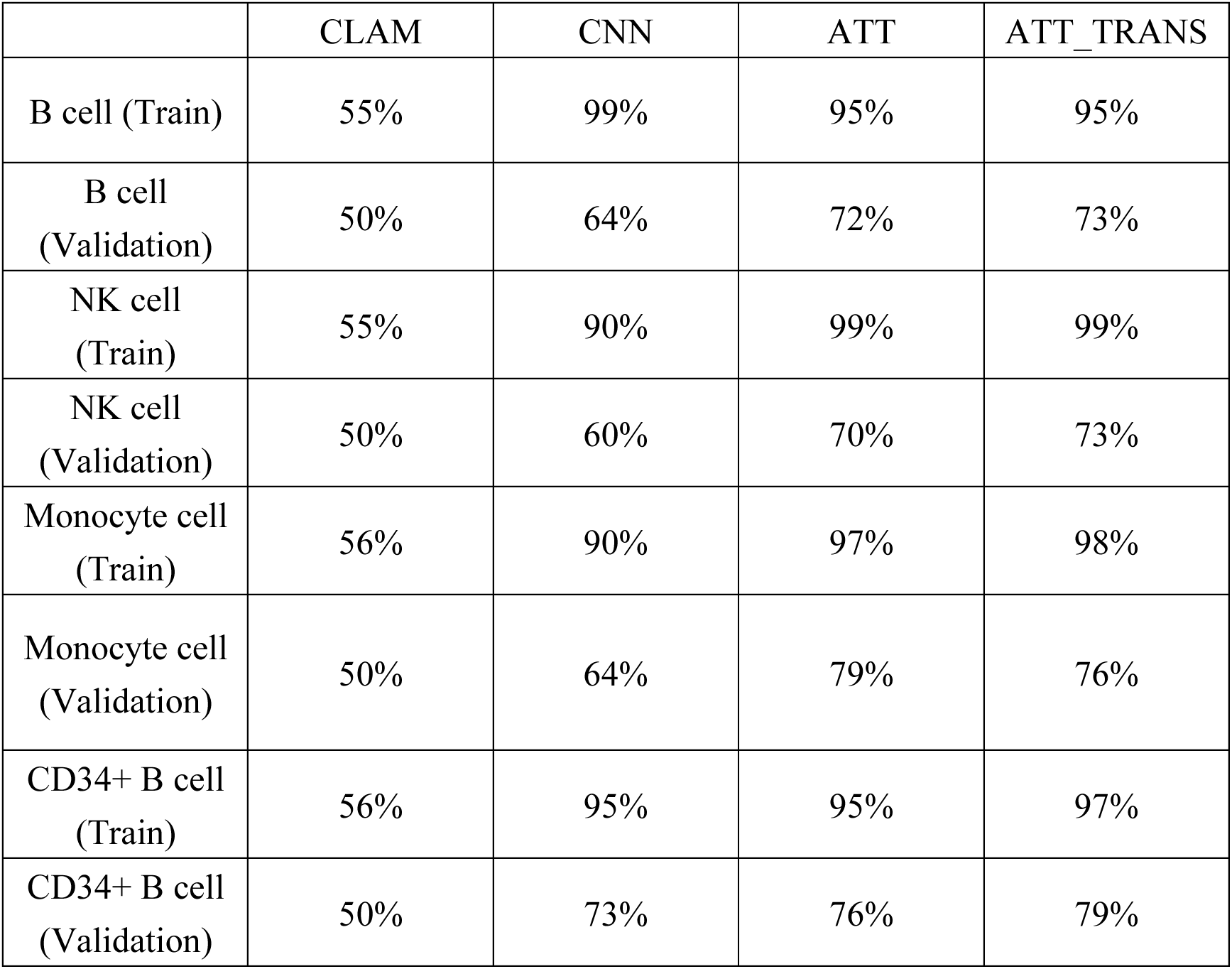
Validation Accuracy of CLAM, CNN, ATT, and ATT_TRANS on B cell, CD34+ B cell, monocyte and NK cell.

|  | CLAM | CNN | ATT | ATT_TRANS |
| --- | --- | --- | --- | --- |
| B cell (Train) | 55% | 99% | 95% | 95% |
| B cell (Validation) | 50% | 64% | 72% | 73% |
| NK cell (Train) | 55% | 90% | 99% | 99% |
| NK cell (Validation) | 50% | 60% | 70% | 73% |
| Monocyte cell (Train) | 56% | 90% | 97% | 98% |
| Monocyte cell (Validation) | 50% | 64% | 79% | 76% |
| CD34+ B cell (Train) | 56% | 95% | 95% | 97% |
| CD34+ B cell (Validation) | 50% | 73% | 76% | 79% |

## Discussion

In this study, we developed LIMPACAT, an integrative framework combining scRNA-seq and digital pathology to accurately infer immune cell compositions associated with prognosis in liver cancer. Using a multiple instance learning (MIL) attention models applied to whole-slide images (WSIs), LIMPACAT successfully predicted immune cell abundance. Our findings specifically identified a positive correlation between the abundance of B cells and NK cells and improved survival outcomes in liver cancer patients. This aligns with previous studies underscoring the roles of B and NK cells in liver cancer immunity. Zhao et al. identified distinct B cell subsets within the hepatocellular carcinoma (HCC) microenvironment, where activated B cells demonstrated heightened metabolic activity, correlating with enhanced effector functions. In contrast, exhausted B cells exhibited diminished functionality, indicating the varied roles of B cell subsets in tumor suppression and immune modulation ^27^. Additionally, Qin et al. reported that high infiltration levels of cytokine-secreting B cells, capable of antigen presentation, were associated with better patient survival rates, further suggesting the immunoregulatory and tumor-suppressive functions of B cells in HCC ^28^. Zou et al. similarly emphasized B cells’ contributions to shaping the immune microenvironment, through their role in T cell activation and antibody production, which collectively promotes antitumor immunity and correlates with favorable patient prognosis ^29^. These observations consistently align with our results, where a greater presence of B cells in the liver tumor microenvironment correlates with improved survival outcomes.

Similarly, NK cells emerged as a crucial component of the immune response in our study, demonstrating significant contributions to antitumor immunity in HCC. This is supported by findings from Xi et al. and Sajid et al., which highlight NK cells’ abilities to mediate cytotoxicity directly through granzyme and perforin release, as well as indirectly via cytokine secretion, such as IFN-γ, which stimulates immune responses against tumor cells ^30,31^. The presence of NK cells was positively correlated with patient survival in our study, echoing these findings and further indicating NK cells’ potential to improve HCC prognosis by enhancing immune surveillance.

Our study employs the attention transformer model within a Multiple Instance Learning (MIL) framework to selectively focus on immune-rich regions in whole-slide images (WSIs), enhancing prognostic predictions by accurately stratifying immune cell levels within the tumor microenvironment. Unlike existing studies, our approach uniquely positions immune cell composition as a primary prognostic factor. For instance, Feng et al. proposed a sliding-attention transformer model to predict T cell receptor and antigen interactions, aiming to facilitate neoantigen discovery for immunotherapy. However, their work focuses on molecular-level immune responses rather than immune stratification in the tumor microenvironment ^32^. Similarly, Duanmu et al. used spatial attention to predict treatment responses in breast cancer pathology, focusing on therapy effectiveness rather than immune cell abundance ^33^.

Additionally, Xiong et al. developed a hierarchical attention-guided MIL framework for WSI classification, which also leverages attention mechanisms to identify critical regions in images. However, their model primarily targets general cancer classification without addressing immune cell stratification, which is central to our study ^34^. Mahmood et al.’s convolution-transformer hybrid model emphasizes adaptive convolution and dynamic attention to capture fine-grained features in renal cell carcinoma images. While effective for classification, it differs from our immune-focused approach tailored for prognostic applications ^35^.

In comparison to Lu et al.’s CLAM model—a weakly supervised MIL framework that applies attention mechanisms for WSI classification—our approach takes a different direction. Although CLAM successfully identifies high-diagnostic-value regions with minimal labeling ^12^, it primarily addresses multiclass pathology classification rather than focusing on immune cell deconvolution for direct prognostic purposes. In our experiments, CLAM reached only 50% accuracy in classifying immune cell levels within the tumor microenvironment, underlining its limitations for this specific task. By contrast, the LIMPACAT model, designed explicitly for immune cell deconvolution, achieved nearly 80% accuracy, marking a substantial improvement in predictive accuracy for patient immune infiltration of B cells and NK cells.

## Conclusion

This study’s LIMPACAT framework (Liver Immune Microenvironment Prediction and Classification Attention Transformer) demonstrates the feasibility of combining scRNA-seq and WSI data to accurately predict immune cell composition and patient prognosis in liver cancer. The findings affirm the importance of comprehensive tumor microenvironment analyses, with B cell, CD34+ B cell, monocyte and NK cell abundance emerging as significant prognostic indicators. Although challenges such as scRNA-seq annotation and model performance. l generalizability remains, ongoing efforts will be essential to refine and enhance this approach. Overall, LIMPACAT contributes critical insights into the cancer immune landscape and introduces innovative strategies for multi-omics data integration and image analysis, underscoring the potential of such models in precision oncology applications.

## Supplementary Figures

Supplementary Figure 1 Correlation matrix showing relationships among filtering metrics (Raw Features, Filtered Cells, Filtered Features, Retained Cell Percentage) in the GSE189903 dataset. Positive correlations confirm the effectiveness of the filtering process

Supplementary Figure 2 Filtered Cell Population with nFeature Counts Between 250 and 2500

Supplementary Figure 3 Distribution of mitochondrial gene percentage (mt percent) and nCount across samples post-filtering. (A) shows the mt percent for each sample, reflecting cell viability and quality, with higher percentages indicating potential cellular stress or damage. (B) displays the nCount distribution, representing the sequencing depth across samples, which captures gene expression patterns specific to different cell types or states

Supplementary Figure 4 Correlation between nCount and mitochondrial gene percentage (mt percent) across samples. The weak correlation indicates that mitochondrial gene content is independent of sequencing depth, suggesting minimal impact of sequencing depth on mitochondrial percentage.

Supplementary Figure 5 Correlation between nCount and nFeature across samples. The strong positive correlation confirms that cells retained after filtering have consistent sequencing depth and gene detection, supporting the reliability of high-quality data for downstream analysis.

Supplementary Figure 6 UMAP clustering of scRNA-seq data post-CCA normalization. (A) UMAP plot showing the distribution of 28 clusters, highlighting additional cell state distinctions. (B) UMAP plot colored by sample identity, revealing the distribution of cells across clusters. Lower ARI score suggests less consistency compared to log normalization.

Supplementary Figure 7 UMAP clustering of scRNA-seq data by sct normalization. (A) UMAP plot with 23 defined clusters, indicating preserved cell type distinctions. (B) UMAP plot by sample identity, showing the spread of cells across clusters. The ARI score indicates moderate clustering consistency and successful batch effect mitigation.

Supplementary Figure 8 nFeature and nCount distributions and sample-to-sample correlations for sct normalization (A, C) and CCA (B, D). The boxplots show the distributions of nFeature and nCount across samples, and the heatmaps display sample-to-sample correlation patterns under each normalization method.

Supplementary Figure 9 Cell type annotations after clustering with sct normalization (A) and CCA (B). UMAP visualizations show identified cell types, based on annotation results following each normalization method.

Supplementary Figure 10 Summary of cell composition deconvolution model performance, normalization comparisons, and survival analysis. (A) Training and validation accuracy/error rates for the cell deconvolution model show no overfitting. (B) ARI comparison of immune cell composition predictions across normalization methods (LOG, CCA, SCT) for LIHC samples. (C) Correlation of immune cell composition and survival times across normalization methods, with CCA and SCT showing consistency. (D) Immune cell composition distributions in LIHC samples, showing similarity between CCA and SCT. (E-H) Survival analysis indicates a positive association between higher levels of B cells, NK cells, and monocytes with longer survival times, whereas a negative association is observed with higher levels of CD34+ B cells.

## Acknowledgements

This work was supported by the grants from the National Science and Technology Council (NSTC).

## Author contributions

Yen-Jung Chiu: Review & Editing, Writing – original draft, Validation, Methodology, Investigation, Formal analysis, Data curation, Supervision, Methodology, Resources.

## Software and data availability

The software is available at https://github.com/holiday01/LIMPACAT.git.

